# Genomic and demographic processes differentially influence genetic variation across the X chromosome

**DOI:** 10.1101/2021.01.31.429027

**Authors:** Daniel J. Cotter, Timothy H. Webster, Melissa A. Wilson

## Abstract

Mutation, recombination, selection, and demography affect genetic variation across the genome. Increased mutation and recombination both lead to increases in genetic diversity in a region-specific manner, while complex demographic patterns shape patterns of diversity on a more global scale. The X chromosome is particularly interesting because it contains several distinct regions that are subject to different combinations and strengths of these processes, notably the pseudoautosomal regions (PARs) and the X-transposed region (XTR). The X chromosome thus can serve as a unique model for studying how genetic and demographic forces act in different contexts to shape patterns of observed variation. Here we investigate diversity, divergence, and linkage disequilibrium in each region of the X chromosome using genomic data from 26 human populations. We find that both diversity and substitution rate are consistently elevated in PAR1 and the XTR compared to the rest of the X chromosome. In contrast, linkage disequilibrium is lowest in PAR1 and highest on the non-recombining X chromosome, with the XTR falling in between, suggesting that the XTR (usually included in the non-recombining X) may need to be considered separately in future studies. We also observed strong population-specific effects on genetic diversity; not only does genetic variation differ on the X and autosomes among populations, but the effects of linked selection on the X relative to autosomes have been shaped by population-specific history. The substantial variation in patterns of variation across these regions provides insight into the unique evolutionary history contained within the X chromosome.

**Significance Statement:** Demography and selection affect the X chromosome differently from non-sex chromosomes. However, the X chromosome can be subdivided into multiple distinct regions that facilitate even more fine-scaled assessment of these processes. Here we study regions of the human X chromosome in 26 populations to find evidence that recombination may be mutagenic in humans and that the X-transposed region may undergo recombination. Further we observe that the effects of selection and demography act differently on the X chromosome relative to the autosomes across human populations. Together, our results highlight profound regional differences across the X chromosome, simultaneously making it an ideal system for exploring the action of evolutionary forces as well as necessitating its careful consideration and treatment in genomic analyses.

## Introduction

Patterns of genetic variation are influenced by factors that vary across genomic regions. Mutation rate (1–5) and recombination rate (6–11) fluctuate across the genome and differ between sexes. Mutation increases diversity by introducing novel variants, and the frequency at which mutations occur, or are removed, contributes to observed patterns of diversity. Recombination affects genetic variation by breaking up regions of linked selection and reducing rates of background selection and genetic hitchhiking (12, 13). In regions where the local recombination rate is high, blocks of linked selection will be broken more efficiently than in regions of low recombination, leading to higher levels of genetic variation. If recombination affects the local mutation rate via double strand breaks, genetic variation will similarly be increased (14).

Some regions of the genome (e.g., the X chromosome, the Y chromosome, and the mitochondria) differ in estimates of genetic diversity due to differences in effective population size (15). Under the infinite sites model, expected nucleotide diversity for diploid organisms is 4*N_e_μ* (16), where *N_e_* is the effective population size and *μ* is the mutation rate. Since diversity is a function of population size, regions of the genome that have a lower relative size are expected to have proportionally lower genetic diversity. The X chromosome, in particular, is composed of distinct regions that differ in *N_e_*. The pseudoautosomal regions exist on both the X and the Y and therefore have a similar *N_e_* to that of the autosomes, while the non-pseudoautosomal regions of the X chromosome exist in two copies in females and one copy in males, effectively resulting in ¾ the effective size of the autosomes. Thus, one would expect differences in genetic diversity across the X chromosome due to differences in *N_e_* across the chromosome.

Selection can also shape patterns of diversity uniquely across the genome. Linked selection reduces diversity in neutral regions that are closely linked to genes (17, 18) and this effect can be more or less pronounced under differing strengths of selection. The effects of background selection and/or genetic hitchhiking are reduced moving away from selected regions, which leads to an expected increase in diversity with increasing distance from these regions. Consistent with this, diversity increases with distance from genes on both the autosomes and X chromosome (19–21). Further, the ratio of X to autosome diversity increases with increasing distance from genes (19–21), suggesting that linked selection is stronger on the X chromosome than the autosomes. This could be because the X chromosome is hemizygous in males, so recessive alleles are thought to be under more efficient selective pressures and linked selection can operate more efficiently in lowering diversity in regions near genes (13, 22–24).

In addition to the processes described above, patterns of human demography strongly affect patterns of genetic variation across populations (25–27). African populations generally have higher genetic diversity compared to non-Africans due to a dispersal event out of Africa that left non-Africans with a subset of African variation (27–29). Further, African populations have significant substructure (30, 31) and deep patterns of demographic history (32) that lead to wide variation in observed diversity. These processes also differentially affect regions of the genome with smaller relative population sizes (e.g., the X chromosome and the mitochondria). The X chromosome, with a smaller effective population size, will be more strongly affected by a bottleneck than the autosomes, but recover faster (33).

As mentioned above, the X chromosome contains several distinct regions that have different evolutionary histories, and which operate under various combinations of the above processes. The sex chromosomes (X and Y) in mammals diverged from a pair of autosomes approximately 180 - 210 million years ago (34). Over time, the X and Y evolved to have different structure and gene content with the Y chromosome losing about 90% of its original genes (35, 36). The differentiation between these two homologs has been theorized to be a result of a handful of distinct inversion events on the Y (37–40) that lead to reduced recombination. Homologous recombination does not occur along much of the length of the X and Y chromosome. However, they share two pseudoautosomal regions (PAR1 and PAR2). PAR1 extends ~2.7 Mb from the tip of the proximal arm of each sex chromosome and facilitates X-Y recombination (35, 40). PAR2 extends 320 kb on the tip of the long arm of each sex chromosome, and evolved independently from PAR1 as a result of at least two X to Y duplication events (41, 42). Recombination rate varies significantly across regions of the X chromosome due to X-Y recombination being constrained to the PARs; PAR1 recombination rate is 20x the genome average (43) and PAR2 recombination rate is ~5x the genome average (44). In addition to the two PARs, there is an X-transposed region (XTR) in humans that was duplicated from X to Y around 3 to 4 million years ago, after human-chimpanzee divergence (35, 45–47). The XTR has undergone a series of inversions and deletions, but it maintains ~98% X-Y homology (35, 48) and contains two genes with functional X-Y homologs (46).

Because of its unique structure, inheritance, and evolutionary history, the X chromosome serves as a unique model for studying how genetic and demographic forces act in different contexts to shape patterns of observed variation. For example, departures from neutral equilibrium expectations of X/A diversity have been used to study sex biases in forces such as migration, admixture, generation time, and reproductive success (15, 19–21, 24, 49–51). In this study we expand on this framework by considering individual regions of the X chromosomes separately and by including a large, global sample of humans (2,504 individuals from 26 different populations sequenced as part of the 1000 Genomes Project (52)) that have experienced a range of different demographic histories. From this data, we calculate measures of diversity, divergence, and linkage disequilibrium to investigate the extent to which linked selection, recombination, mutation rate, and demography shape relative patterns of variation across the human X chromosome. This design allows us to better understand the forces that shape genetic variation and gives an unprecedented look into the evolutionary biology of the sex chromosomes.

## Results

### Genetic variation is consistently elevated in the PARs and XTR across human populations

We measured nucleotide diversity in 26 human populations (Table S1) from The 1000 Genomes Project (52) and observed substantial variation across regions of the X chromosome (Figure 1A). Overall, we found that diversity is significantly higher in both PAR1 and XTR than chrX (which we define here as non-pseudoautosomal sequence on the X chromosome not in PAR1, PAR2, or XTR) in nearly all populations (Table S2). We also observed higher diversity in PAR2 than chrX in all cases, but the difference was never significant. After filtering, PAR2 has approximately 15% as many variant sites as PAR1 and approximately 65 kb of callable sequence (Table S3). Due to its size, unusual evolutionary history, and the small amount of data available after filtering, we report observational results for PAR2 but exclude it from interpretations.

**Figure 1.**
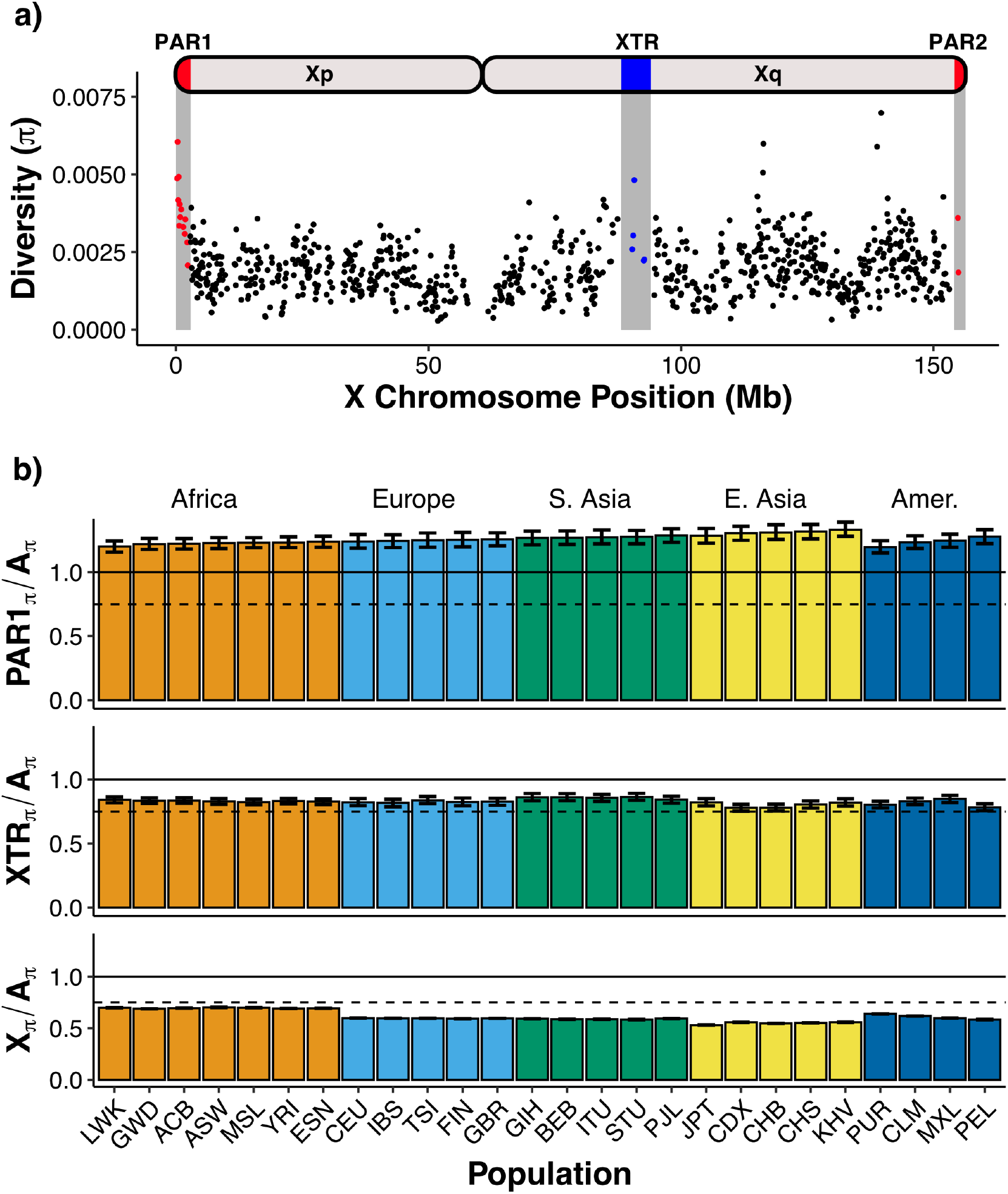
Genetic diversity across regions of the human X chromosome. **A)** Nucleotide diversity is calculated in non-overlapping 100kb windows across the X chromosome corrected for mutation rate variation using hg19-canFam3 divergence. Red indicates the pseudoautosomal regions (PAR1, PAR2) and blue indicates the X-transposed region (XTR). Diversity is calculated using all 1000 Genomes phase 3 samples. **B)** Genetic diversity is calculated in each population between each region of the X chromosome—pseudoautosomal region 1 (PAR1), X-transposed region (XTR) and chromosome X—and the autosomes (chr8). Autosomal and X-linked diversity are corrected for mutation rate (hg19-canFam3 divergence). The solid line at 1.0 represents the null expectation of four PAR1 regions for every four autosomes. The dashed line at 0.75 represents the null expectation of three X chromosomes for every four autosomes. Error bars represent 95% bootstrapped confidence intervals using 1000 replicates. Populations are organized by superpopulations. Individual population abbreviations are labeled, and full names are available in Table S1.

### Ratios of PAR/A and chrX/A diversity exhibit opposite patterns across human populations

To further explore differences in diversity among regions on the X chromosome, we divided diversity values calculated in PAR1, XTR, and chrX by those from chr8 (referred to as autosome or A), for each of the 26 populations (Figure 1B). chrX/A values were below the neutral expectation of 0.75 (assuming equal sex ratios and 3 X chromosomes for every 4 autosomes). PAR1/A ratios were all greater than 1.0, and thus greater than expectations based on chromosome counts (i.e., two copies of chromosome 8 in all individuals, and two copies of PAR1, either on two X chromosomes in females or on the X and Y in males). We observed PAR1/A ratios around 1.25 within Africa, and gradually increasing PAR1/A ratios in populations outside of Africa. In contrast, chrX/A ratios decreased in populations outside of Africa. This pattern was recapitulated in admixed American populations, in which we observed that PAR1/A ratios increased with decreasing African ancestry proportions, while chrX/A ratios decreased with decreasing proportion of African ancestry. In order of decreasing African ancestry proportion, those populations are Puerto Rican (~28% African ancestry), Colombian (~7%), Mexican (~4-5%), and Peruvian (~2%) (53–57).

Surprisingly, our observations of XTR did not match our expectation that the XTR and chrX would behave similarly because both are present in one copy in genetic males and two copies in genetic females. Across populations, XTR/A diversity was consistently greater than observed chrX/A diversity (Figure 1B) and variation in XTR/A ratios did not appear to correspond with demography.

### Substitution rate varies across regions of the human X chromosome

Mutation rate, which is known to vary across the genome (58–60), influences observed levels of genetic diversity. Under a neutral model of evolution, higher mutation rates result in more genetic variation and thus increased levels of diversity (61). To explore regional variation in mutation, we used substitution rate (divergence) between the human and dog reference genomes as a proxy. In general, divergence did not increase with increasing distance from genes (Figure 2), though the XTR exhibited a slightly elevated substitution rate in the bin filtering 20kb from genes and PAR2 substitution rate fluctuated with increasing distance from genes (Table S3).

**Figure 2.**
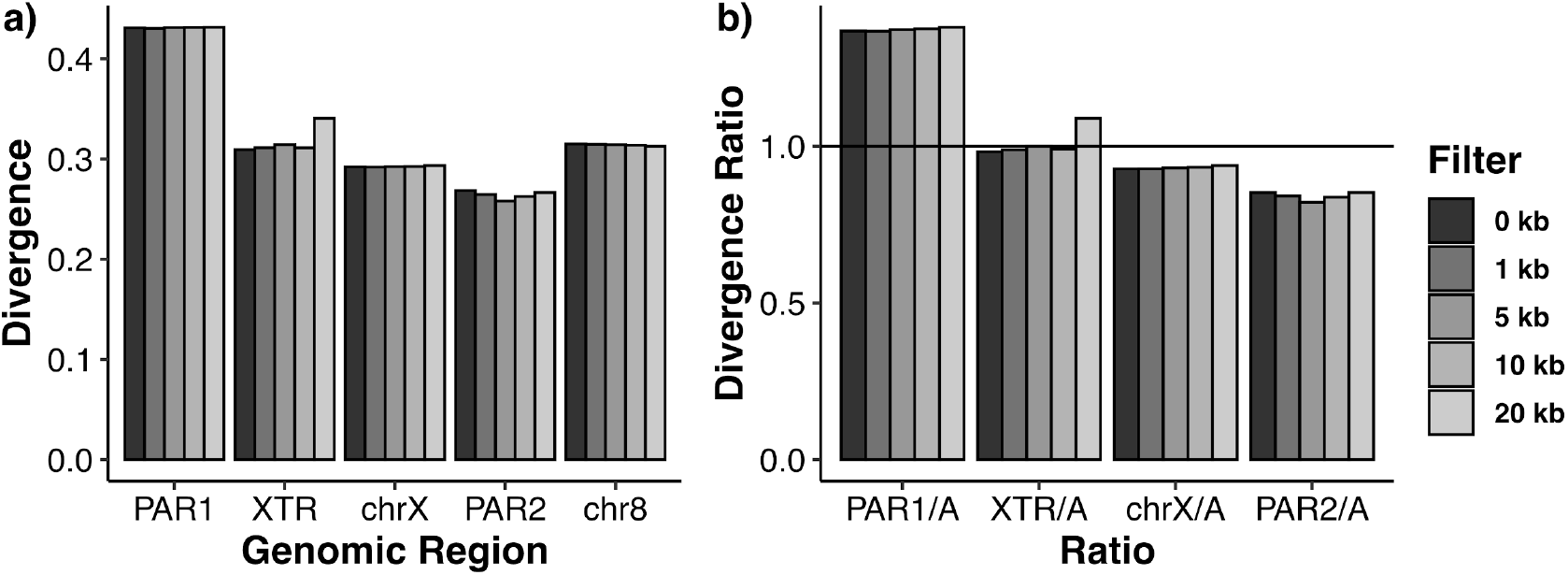
Divergence across the genome. Human-dog divergence (hg19-canFam3) **A)** across regions of the human X chromosome and chromosome 8; and **B)** between each region of the human X chromosome relative to chromosome 8. A solid horizontal line is placed at a divergence ratio of 1, which would imply an equal mutation rate between regions. Divergence is computed for all intergenic regions with no filter with distance from genes (0kb), or with filtering out regions near genes (1kb, 5kb, 10kb or 20kb). The number of base pairs in each region is reported in Table S3.

While we did not find an association between human-dog divergence and distance from genes, we observed striking differences in substitution rates across the different regions of the X chromosome and chromosome 8 (Figure 2). PAR1 had the highest substitution rate (~1.3x that of chr8), while the substitution rate of the XTR was similar to that of chromosome 8. Both chrX and PAR2 had lower substitution rates than chromosome 8. For chrX, the difference was slight (0.93x that of chr8).

### Linkage disequilibrium in PAR1 and XTR is lower than the rest of the X chromosome

We calculated average *r^2^* across the X chromosome and chromosome 8 to characterize linkage disequilibrium (LD) as a proxy for recombination rate (see Methods). Consistent with our expectation that a higher recombination rate will break up linkage, we found that LD is lowest in PAR1 and highest in chrX, with chr8 exhibiting values slightly lower than chrX (Figure 3). However, the XTR exhibited intermediate *r^2^* values that fell approximately halfway between PAR1 and chrX (22 of 26 populations, Figure S1). Estimates for LD in PAR1 and the XTR varied slightly between populations within the same superpopulation (Figure S1), but these trends are broadly consistent across all populations studied.

**Figure 3.**
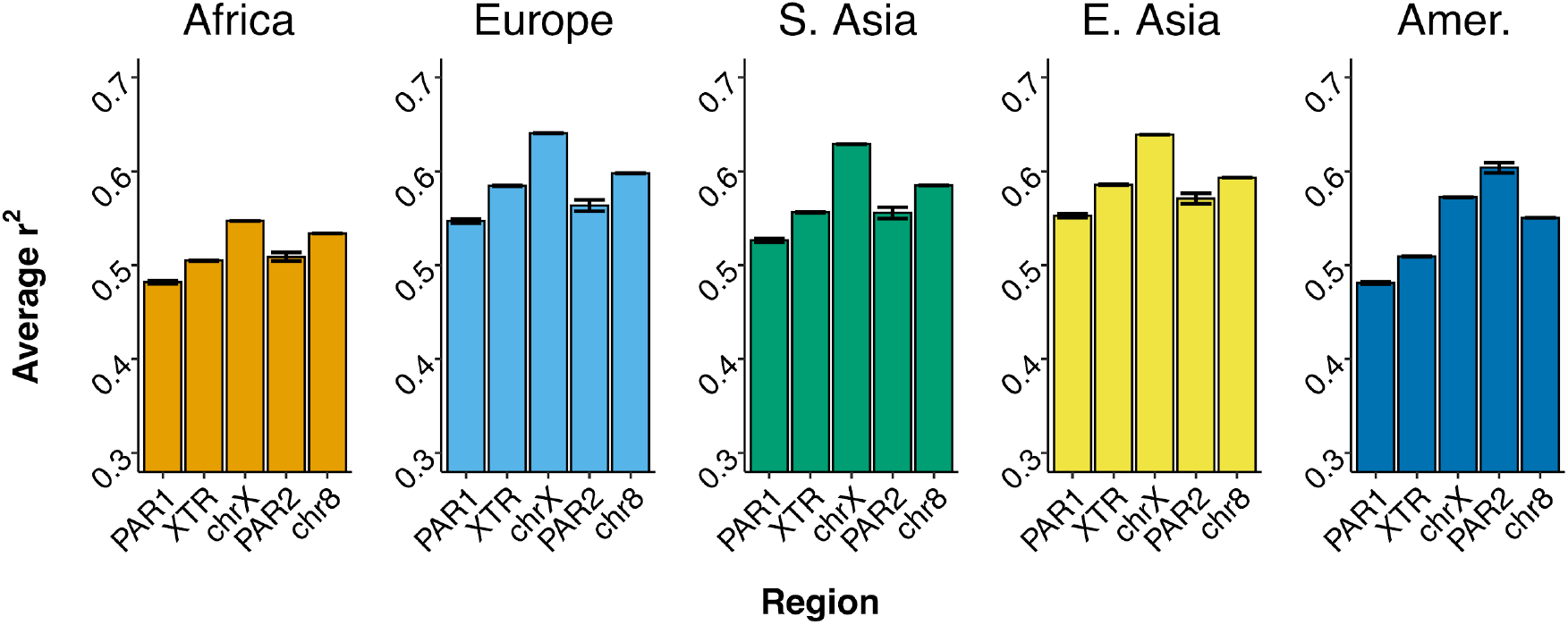
Average linkage disequilibrium across genomic regions. Linkage disequilibrium (LD) is calculated in each X chromosome region and chromosome 8 for each superpopulation. LD is calculated for each site in a given genomic region by averaging all pairwise *r^2^* values +/-300kb from that site. Average *r^2^* values for each site are then used to calculate mean LD for a given region. Error bars represent 95% bootstrapped confidence intervals (1000 replicates).

### Genetic diversity and linkage disequilibrium are negatively correlated on the X chromosome

We used a linear regression analysis to explore the relationship between LD and diversity in the same windows. We found a significant negative relationship between LD and diversity (R^2^ = 0.127, P = 2.46×10^-31^; Figure S2A). We also observed that LD explains less variation in diversity as we filtered further from genes (R^2^ = 0.10, P = 8.03×10^-22^; Figure S2B).

### Regions of the X chromosome exhibit contrasting and population-specific patterns of linked selection

We explored how X/A, PAR/A, and XTR/A diversity ratios varied across 26 populations in Africa, Europe, South Asia, East Asia, and the Americas as we filtered out regions close to genes (Figure 4A). We calculated diversity in each population after filtering out genes and conserved sequences. We then iteratively filtered out regions of increasing size, starting from the region closest to genes and moving out to a designated threshold (1kb, 5kb, 10kb, 20kb). The measure we use is the difference between (a) diversity calculated after filtering flanking sequence from genes and (b) diversity calculated where we only filter genes. More efficient selection on the X chromosome should lead to patterns of increasing X/A diversity ratios moving away from genes due to more pronounced linked selection on the X chromosome (19–21). Consistent with this prediction, we found that X/A diversity increased as we filtered farther from genes in four of the five superpopulations (Figure 4A). In contrast, we surprisingly found that PAR/A and XTR/A diversity ratios decreased as we filtered out regions close to genes.

**Figure 4.**
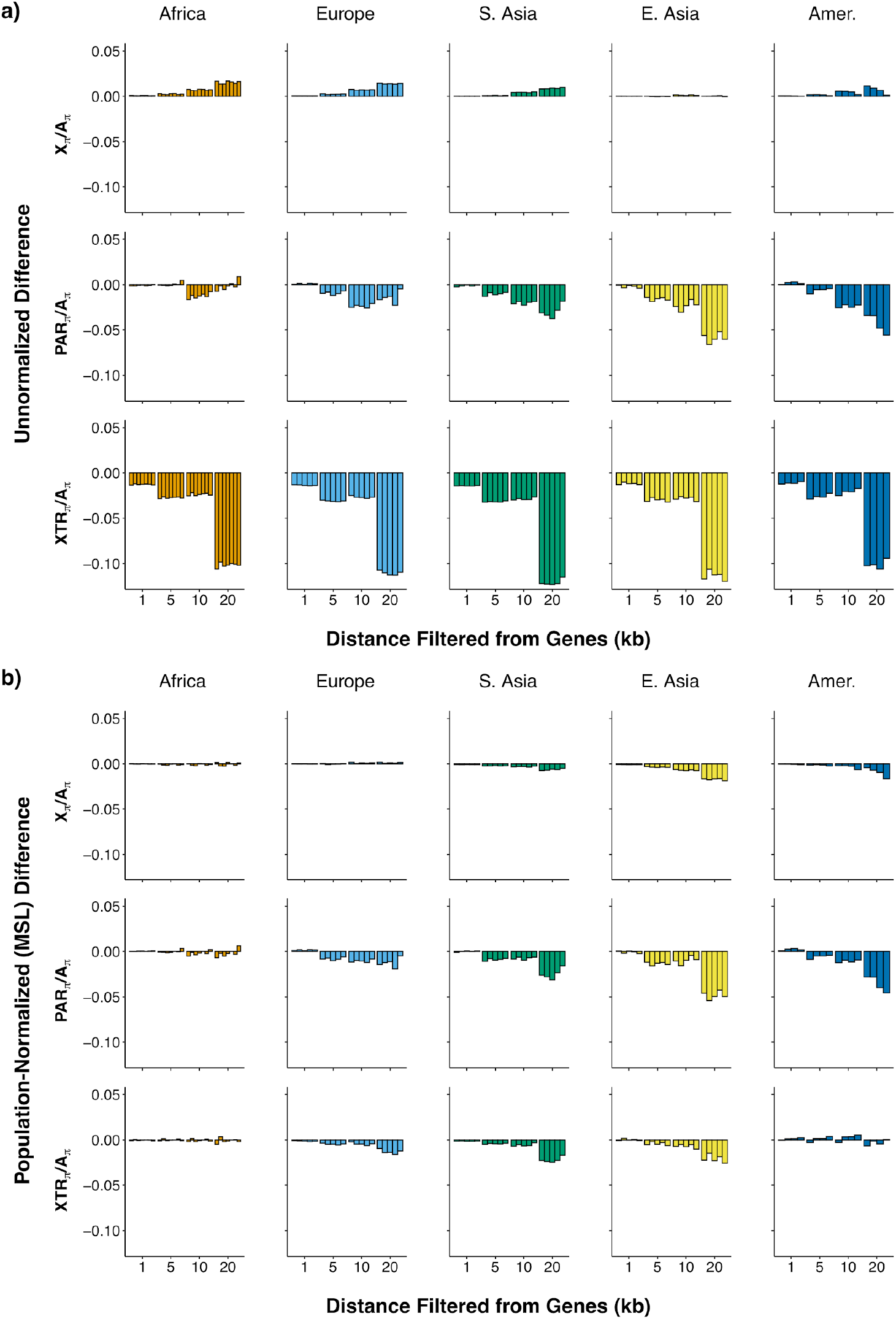
Ratio of X to autosomal diversity with increasing distance from genes across populations. **A)** Diversity ratios are reported between regions on the X—non-pseudoautosomal X (X), pseudoautosomal region 1 (PAR), and X-transposed region (XTR)—and the autosomes for 26 populations from the 1000 Genomes Project. Values are reported as the difference between using a filter for only genes and a filter for 1kb, 5kb, 10kb, and 20kb from genes, respectively. The order of populations is the same as reported in Figure 1B. **B)** These ratios are demography normalized by reporting each population relative to Mende in Sierra Leone (the population with the highest nucleotide diversity among all populations for most X-chromosomal regions)

To account for the effect that demography may have on these patterns, we corrected the ratios for each of the above populations by dividing these ratios by those from the African population MSL (Mende in Sierra Leone)—the population in our dataset exhibiting the greatest diversity across most regions of the X chromosome (Figure 4B). If patterns of linked selection are consistent across populations, we expect the normalized differences to be equal to 0 across all regions with increasing distance from genes (21). Overall, we found that while most populations had normalized X/A ratios that did not change with distance from genes, some non-African populations, particularly in East Asia, displayed values that decreased with distance from genes (Figure 4B).

For PAR/A and XTR/A ratios, the effect of normalizing to MSL was less pronounced (Figure 4B). Although there is no effect of distance from genes on the normalized ratio in African populations, the PAR/A ratio still decreases in non-African populations. XTR/A ratios have their effect of distance from genes reduced after normalization, but there is still some decrease evident in non-African populations.

To look at the effect of different outgroup populations on normalizing X/A ratios, we divided diversity ratios for the 26 populations by the populations with the highest diversity in each of the four remaining superpopulations (Figure S3A–D). When we normalized the 26 populations by the Tuscan population (TSI; the highest diversity population in Europe for most of the X-chromosomal regions), we found that X/A, PAR/A, and XTR/A ratios in the remaining European populations were unaffected by increasing distance from genes with PAR/A and XTR/A ratios fluctuating across populations that were members of the other four superpopulations (Figure S3A). We repeated this normalization for each of the three remaining superpopulations (Figure S3B–D) and each one exhibited the same pattern described above: X/A, PAR/A, and XTR/A ratios were unaffected by distance from genes for populations that were part of the superpopulation used for the normalization, while PAR/A and XTR/A ratios varied among other populations.

## Discussion

There are many processes that shape the landscape of genetic variation across the genome. Patterns of genetic variation across the X chromosome are especially complex because its unique structure and pattern of inheritance have the potential to interact with these processes in different ways. In this study, we examined 26 diverse human populations and found remarkable variation in genetic diversity on the X chromosome, both among populations and across different regions of the X chromosome itself. More specifically, we found that the landscape of genetic variation across the X chromosome was structured by mutation, recombination, and population history, which differentially affected major regions of the X chromosome—the PAR, XTR, and nonPAR—and led to substantial variation in genetic diversity across these regions.

### The X-transposed region has intermediate properties of both PAR1 and nonPAR

Of the X-chromosomal regions we studied, our results for the X-transposed region (XTR) were especially surprising because its properties were intermediate to both the pseudoautosomal regions and the nonPAR regions of the X chromosome. Though the XTR shares homology with the Y chromosome, we expected it to behave similarly to the nonPAR regions of the X chromosome because it underwent an inversion preventing recombination (48). However, in our measures of diversity (Figure 1B) and recombination (Figure 3), the XTR exhibited values that were greater than we observed in nonPAR, but less than we observed in PAR1.

The unusual pattern of diversity within the XTR could be driven, in part, by technical artifacts. We recently showed that, due to homology, the X-transposed sequences between the X and the Y are similar enough to confound the mapping of raw sequencing reads (62). This leads to lower mapping quality and sequencing depth, which in turn reduces the number of variants called (62). With this in mind, our observations of higher genetic diversity in the XTR than the nonPAR in this study are still surprising, as they are likely lower than they should be because the sex chromosome data from the 1000 Genomes Project were not corrected for this problem, having been published before the problem and correction were described by Webster et al. (2019) (62).

Our observation of lower linkage disequilibrium (LD) in XTR, if LD serves as a good proxy for recombination in this case, is consistent with recombination occurring in this region. This is unexpected because, despite its X-Y homology, the XTR experienced an inversion event that is proposed to have prevented further recombination from occurring (48). Our results are instead consistent with more recent work suggesting that there is evidence for unequal crossing over (63) in the XTR for a small portion of the population, leading some researchers to dub this region PAR3 (47). Though it remains unlikely that the X-Y recombination in this region is extensive, the substantial difference between XTR and nonPAR that we observed in this study should motivate further molecular investigations of the XTR to better understand this behavior.

Our results also have methodological implications. Building on our previous description of technical artifacts in this region (62) and in accordance with previous observations about X-transposed region diversity (64), the elevated divergence and recombination that we observe in the XTR further establish that this region needs to be removed from analyses of the nonPAR X chromosome. Many genome-wide studies remove the PARs when analyzing the X chromosome, but we suggest that it is equally important to remove the XTR.

### Recombination influences mutation rate across X-chromosomal regions

While past studies have generally found that genetic divergence is not associated with recombination hotspots across the autosomes (65), we observed a correlation between our proxies for recombination rate and mutation rate on the X chromosome when considering the X-chromosomal regions. Our substitution rate observations (Figure 2) are consistent with mutation rate being higher in PAR1 and XTR than the nonPAR regions of the X chromosome. Similarly, our linkage disequilibrium estimates (Figure 3) are consistent with higher recombination in PAR1 and XTR than nonPAR. PAR1 has been previously observed to have increased substitution rate relative to autosomes (66); a result which we have also confirmed here. This phenomenon supports the conclusion that recombination rate is positively correlated with mutation rate. Further, this pattern is replicated in 22 of 26 populations (Figure S1), consistent with it having a more general biological explanation, rather than a demographic one.

Specifically, these observations support the hypothesis that recombination can be mutagenic in humans. Results from *S. cerevisiae* have demonstrated that double-strand-break repair can be mutagenic (67). Additionally, it has been argued that the correlation between recombination rate and genetic diversity in the human PAR1 is driven by the relationship between recombination rate and divergence (68). Here our observations expand this observation to the other regions of the X chromosome and suggest that recombination influences mutation rate in a region-specific manner, especially in regions with extraordinarily high recombination rates (like PAR1).

### The X chromosome exhibits patterns of linked selection that differ among populations

The ratio of X chromosome to autosome diversity has long been of interest in exploring aspects of population history, particularly those that are sex-biased (15, 19, 21, 51, 69). Analyzing these ratios in the same way across 26 human populations gives us an unprecedented look at how this measure changes across a variety of demographic histories and different regions of the X chromosome.

When considering just the nonPAR X, we observe a pattern largely in line with previous studies: the highest ratios are in African populations (Figure 1B). As population bottlenecks disproportionately affect the X chromosome because of its smaller effective population size (33), lower ratios outside of Africa were likely the result of a bottleneck in the population ancestral to all non-African groups when it was migrating out of Africa (28). Other work has shown that strong male biases during this migration might have also decreased these ratios (70, 71). Interestingly, when we organized admixed populations from the Americas based on the amount of African ancestry they contain, we recapitulated the same pattern: we observed decreasing X/A ratios with decreasing African ancestry. Thus, while there is clearly variation among individual populations, the migration out of Africa by some groups is by far the most dominant force shaping X/A ratios in humans.

In contrast to the nonPAR X, when we used XTR/A or PAR1/A ratios, we observed very different patterns (Figure 1B). Both ratios were significantly higher than expected, with XTR/A ratios greater than 0.75 and PAR1/A ratios greater than 1.0 in all populations. Moreover, for PAR1/A ratios, we observe an inverse demographic pattern to what we observed for X/A, with PAR1/A ratios increasing out of Africa and admixed American populations exhibiting increasing values with decreasing African ancestry. It is critical to note that the only difference among these analyses is the region being studied, so that PAR1, XTR, and the nonPAR X display these contrasting patterns within the same populations and under the same demographic histories. For the XTR, ratios higher than both the nonPAR X and a neutral expectation of 0.75 could be consistent with some recombination in this region, as discussed above, but it’s unclear why differences among populations don’t scale with those observed in nonPAR X and PAR1. Mutagenic recombination might explain the higher than expected PAR1 values overall, but it does not immediately explain the apparent increase in PAR1/A ratios out of Africa.

We explored the trend by simulating two regions with different recombination rates under the out-of-Africa model of human evolution. We observe that the interaction of differences in recombination (e.g., between PAR1 and nonPAR X) and differences in population history between different human groups (e.g., European vs. African populations) can explain both the higher than expected PAR1/A ratios and the increase in these ratios in non-African populations (Figure S4). Another possibility, which we do not explore here, is the effect of balancing selection, which could maintain greater genetic diversity in PAR1 relative to the autosomes if it were acting disproportionately in this region (72, 73).

When considering the effect of linked selection, we saw that nonPAR X/A ratios increase after filtering farther from genes which is consistent with the hypothesis that the X chromosome experiences more efficient diversity-reducing selection (i.e., hitchhiking and background selection) than the autosomes, due in part to it being found in only one copy in most genetic males (19, 21). In contrast, we observed decreasing PAR/A and XTR/A ratios with distance from genes (Figure 4A) which could be consistent with multiple processes including less efficient selection and a higher density of elements under strong selection in these regions than the autosomes.

In order to learn more about the differences in the X/A, PAR/A, and XTR/A ratios across the populations we studied (Figure 1B), we considered relative ratios between sets of two populations (“normalized ratios”; Figure 4B). Previously, Arbiza et al (2014) plotted these normalized ratios as a function of distance from genes in order to separate the effects of demography and selection (21). If normalized ratios don’t change with distance from genes, it implies that demography drives any observed differences in X/A diversity ratios among populations. However, If these ratios do change with increasing distance from genes, it suggests that population differences in patterns of selection can be shaping X/A diversity ratios as well.

Arbiza et al. (2014) found that, for two populations (from EUR and EAS), this normalization resulted in roughly equal X/A ratios near and far from genes, leading authors to conclude that the general relationship of selection between the X and autosomes was similar across human populations (21). We replicated this normalization for PAR/A, XTR/A, and X/A ratios (using Mende in Sierra Leone, MSL, as our denominator for all populations) and observed the same result of Arbiza et al. (2014): no increase in X/A ratios with increasing distance from genes in African and European populations (Figure 4B). However, when we normalized using populations with vastly different demographic histories, we found some notable differences in X/A ratios which suggest that selective forces vary across human populations. First, PAR/A and XTR/A ratios always increased in populations outside of the superpopulation used for the normalization. Second, each ratio was unaffected by distance from genes when it was normalized by its own superpopulation. While there may be common effects of selection on the X chromosome in some populations, it is likely that different regions are under different selective pressures across different global populations.

This builds on a growing picture that shows if we are to fully understand genomic variation and human evolutionary history, we need to look at a diversity of populations (74). While normalization provided a simple, straightforward picture when considering two human superpopulations (21), the inclusion of additional populations demonstrated that this picture is far more complex and requires more nuanced interpretations. Many interpretations in population genetic studies depend on the choice of the populations that are being compared (75). When making genomic claims, we must carefully consider the context of the populations that we are comparing. Further, analyses that include multiple genomic regions can shed light on how evolution shapes the genome as a whole. In this study, without studying many populations from around the world we would not have been able to differentiate phenomena that seem to be shared across humans (e.g., biology of the XTR) versus those that vary among groups (e.g., patterns of linked selection on the X). Importantly, the X chromosome is an important region for teasing apart these global and populations-specific evolutionary processes.

## Methods

### Human DNA variation data

We obtained human genetic variant data in the form of VCF files from Phase 3 of The 1000 Genomes Project from hg19 (52). We analyzed data from the X chromosome and chromosome 8 (chr8), an autosome approximately the same length as the X chromosome, in 26 different populations from 5 major geographical regions (broadly, Africa, Europe, South Asia, East Asia, and the Americas; Table S1). Throughout this paper, we use “superpopulation” to refer to the grouping of all individuals within a major geographical region (e.g., the superpopulation “Africa” refers to samples from all populations in Africa) and “population” to refer to one of the local populations (n=26). We used the strict mask provided by The 1000 Genomes Project (*20141020.strict_mask.whole_genome.bed*) to assess callability and determine the number of monomorphic (i.e., invariant) sites in each region.

### Filtering regions of the genome

We used the UCSC Table Browser (76) to obtain coordinates for genomic elements that may be affected directly by selection or difficult to align. We obtained coordinates for whole genes (transcription start to transcription end), centromeres, telomeres, CpG islands, and simple repeats. To curate a comprehensive and conservative list of whole genes we intersected records from the RefSeq genes track, the GENCODE genes track, and the UCSC genes track. We created iterations of this record with 0kb, 1kb, 5kb, 10kb, 20kb, 50kb, and 100kb of flanking sequence upstream and downstream of each gene record additionally filtered out. For our main analyses, we used the filter that excluded 10kb flanking whole genes to better control for linked selection. We chose to filter 10kb from genes because filtering with greater distance from genes resulted in removing much of the sequence from our regions of interest on the X chromosome (Table S3). We processed all filter coordinates using bedtools (77).

### Divergence

To account for mutation rate variation, we corrected our diversity estimates in each region using pairwise divergence values between human and dog (hg19-canFam3) reference genomes. We obtained divergence estimates for each filter and window type by applying the Estimate Substitution Rate tool to sequence alignments from the Galaxy Toolbox (78) and correcting these results using the Jukes-Cantor 1969 model (79).

In addition to using divergence estimates to account for variation in mutation rate, we explored how hg19-canFam3 divergence estimates within PAR1, PAR2, XTR, chrX, and chr8 change as we filter with increasing distance from genes (0kb, 1kb, 5kb, 10kb, 20kb). We applied the same filters for regions affected by selection used for diversity analysis (see below). We additionally calculated divergence ratios between each of the X chromosome regions relative to chromosome 8 (Figure 2).

### Diversity calculations

We estimated uncorrected and unnormalized genetic diversity as the average number of pairwise differences per site (π) among sequences in each population. We used allele frequencies of single nucleotide polymorphisms to calculate diversity for each variant site:

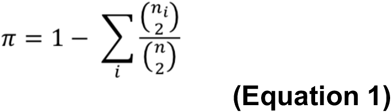

where *n_i_* is the allele count of allele *i* in a sample and *n* is the sum of *n_i_* (80). We calculated diversity across chrX and chr8 for each of the 26 1000 Genomes populations (Table S1) in 1) non-overlapping 100 kilobase (kb) windows partitioned across each analyzed chromosome, and 2) distinct regions across the X chromosome: the pseudoautosomal regions located at the tips (PAR1 and PAR2), the X-transposed region located on the long arm (XTR), and the remaining regions (referred to simply as chrX; see Figure 1A). We obtained coordinates for PAR1 and PAR2 from build hg19 of the human genome and coordinates of XTR from (35). See Table S4 for coordinates. For chromosome 8, we performed all calculations across the entire chromosome.

In each window/region we corrected for differences in mutation rate by dividing the window by the corresponding calculation of substitution rate in that window/region. After correcting for divergence in each of our 100kb windows, we used permutation tests to compare mean diversity among the X chromosome regions for each of the 26 1000 Genomes populations. We divided chrX into 100kb non-overlapping windows and we permuted these windows 10,000 times to test the significance of the difference between each X chromosome region (PAR1, XTR, PAR2) and the rest of chrX.

### Diversity ratios between the X and autosomes

To explore variation across the X chromosome regions (PAR1, XTR, chrX), we calculated the ratio of diversity corrected for divergence in each region relative to diversity corrected for divergence on chr8. We did this for each human population. We calculated 95% bootstrapped confidence intervals (1000 replicates with resampling) for each ratio.

### Normalizing diversity for human demography

To explore the role that demography plays across these regions, we normalized diversity on the X and autosome, both corrected for divergence, by dividing by the population with the highest level of estimated diversity (in this case Mende in Sierra Leone; MSL). Thus, we have estimates of normalized diversity for 25 of the 1000 Genomes populations.

### Effects of linked selection on unnormalized and normalized diversity

To explore the effects that linked selection has on diversity, we analyzed sequence diversity with increasing distance from genes (0kb, 1kb, 5kb, 10kb, 20kb). To visualize the effects of filtering potentially linked sequences, we plotted the difference in diversity between each filter that removed flanking regions from genes (1kb, 5kb, 10kb, and 20kb) with the measurement of diversity that only excluded genes and no flanking sequence (0kb). We did this both for unnormalized diversity and for measurements of diversity normalized to MSL (Figure 4B) as well as TSI, PJL, KHV, and PUR (Figure S3A–D).

### Linkage Disequilibrium

We used linkage disequilibrium as a proxy to explore recombination rate variation across the X chromosome and chromosome 8. We first applied the same filters discussed above, filtering 10kb of sequence flanking genes. We then calculated average *r^2^* in 100 kb windows across each chromosome as well as within the X chromosome regions and all of chromosome 8. We did this separately for each superpopulation (Figure 3) and for each of the 26 1000 Genomes populations (Figure S1). We considered each site individually and averaged all of the pairwise *r^2^* values (calculated with Plink (81)) between that site and all other sites within 300kb in either direction. We then took the mean of each site’s average *r^2^* values within each 100kb window and within each of our genomic regions (the X chromosome regions and chromosome 8). We estimated 95% bootstrapped confidence intervals by resampling 1000 times in each region of interest. To explore the relationship between LD and diversity, we used linear regression analysis to compare the average *r^2^* values and diversity values calculated in 100kb windows across the X chromosome (Figure S2).

### Simulations

In order to explore the effect of recombination on neutral genetic diversity, we simulated two regions with different recombination rates using SLiM version 3.4 (82). We simulated these regions under the Gravel et al. (2011) model of human evolution (83). We defined two unlinked regions: one with a recombination rate of 2×10^-7^ and the other with a recombination rate of 1×10^-8^. We simulated neutral mutations with a uniform mutation rate of 2.36×10^-8^. Starting in generation 55,960 we took 250 samples (with 100 individuals each) from the African population and European population every 20 generations. We used these samples to construct a bootstrapped distribution for genetic diversity (calculated at each site using Equation 1 and averaged) in each region and plotted this trajectory over time. We did this for each region separately (Figure S4A&B) as well as for the ratio of the two regions (Figure S4C). We tracked the mean value of the distribution as well as the 95% confidence intervals.

## Data Access

We retrieved all data from the publicly available repository at The 1000 Genomes Project (www.internationalgenome.org/data/). We host all code used to analyze the data and generate the figures in this manuscript, which is implemented in a Snakemake pipeline (84), on GitHub (github.com/djcotter/chrX_regional_variation).

## Acknowledgements

We thank Alan Rogers and Jazlyn Mooney for helpful discussion. We acknowledge Research Computing at Arizona State University for providing high-performance computing and storage resources that have contributed to the research results reported within this paper (URL: http://www.researchcomputing.asu.edu). This work was supported by the National Institute of General Medical Sciences (NIGMS) of the National Institutes of Health (NIH) grant R35GM124827 to MAW and a National Science Foundation (NSF) Graduate Research Fellowship to DJC.

## Author Contributions

MAW conceived the research; MAW, THW, and DJC designed the research; DJC and THW analyzed the data; DJC wrote the manuscript with input from THW and MAW; DJC, THW, and MAW edited and approved the manuscript.

