## Supplemental Material for "Genomic and demographic processes differentially influence genetic variation across the X chromosome"

#### Title



#### Classification

Biological Sciences: Genetics

#### Keywords

X chromosome, pseudoautosomal region, X-transposed region, demography, diversity, divergence

### Supplementary Material

#### Supplementary Figures

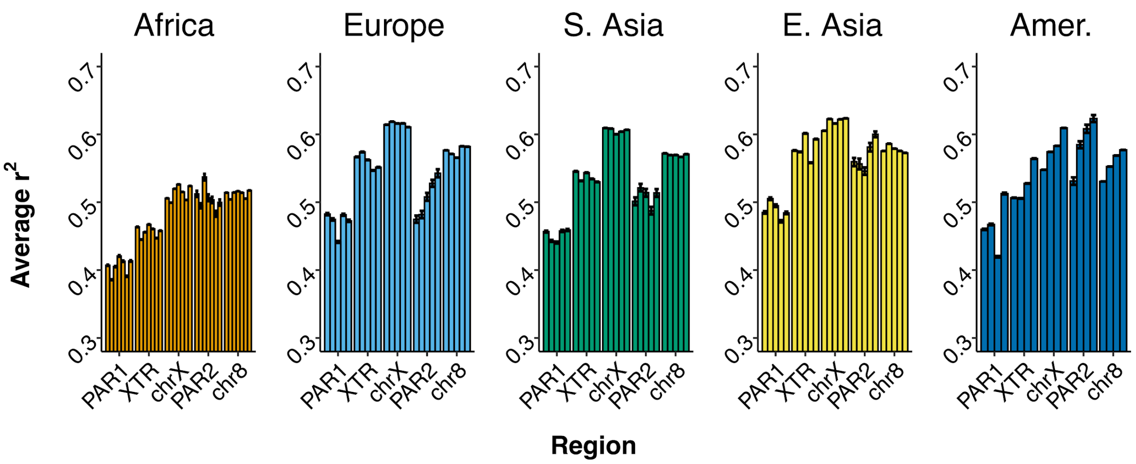


**Figure S1. Average linkage disequilibrium across genomic regions**. Linkage disequilibrium (LD) is calculated in each X chromosome region and for chromosome 8 for each 1000 Genomes Population. LD is calculated for each site in a given genomic region by averaging all pairwise $r^2$ values +/- 300kb from that site. Average $r^2$ values for each site are then used to calculate mean LD for a given region. Error bars represent 95% bootstrapped confidence intervals (1000 replicates with replacement).

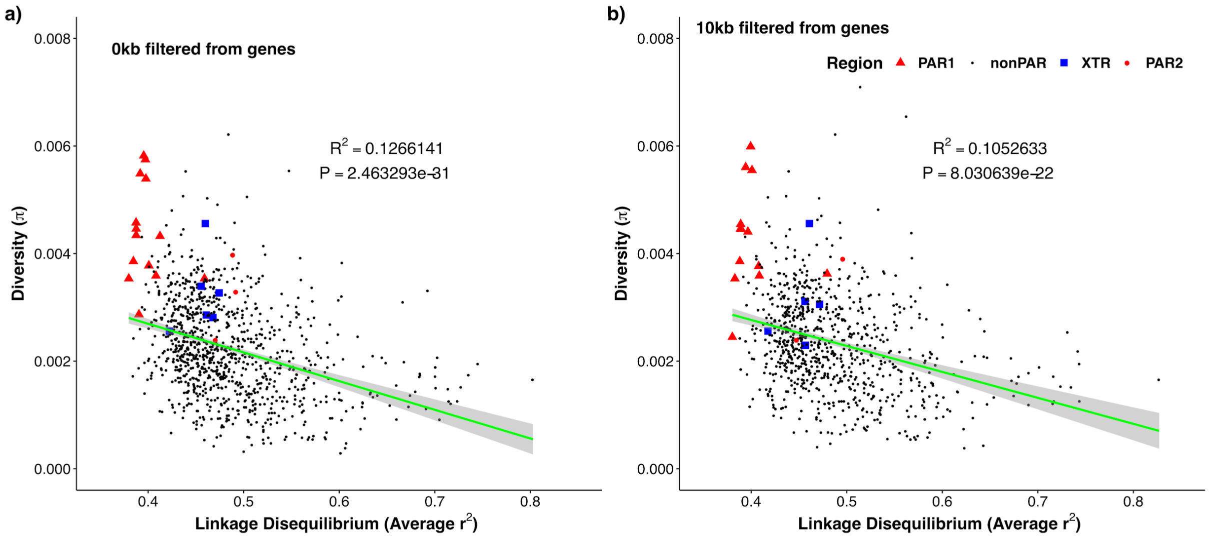


**Figure S2. Linkage disequilibrium and nucleotide diversity across the X chromosome.** Average linkage disequilibrium was calculated in 100kb windows and plotted against corresponding average nucleotide diversity in 100kb windows (corrected for mutation rate with hg19-canFam3 divergence). This was done for **A)** diversity calculated by only filtering for genes and **B)** diversity calculated by filtering for genes +/- 10 kb flanking regions. $R^2$ values for the negative correlation are reported on each plot.

a)

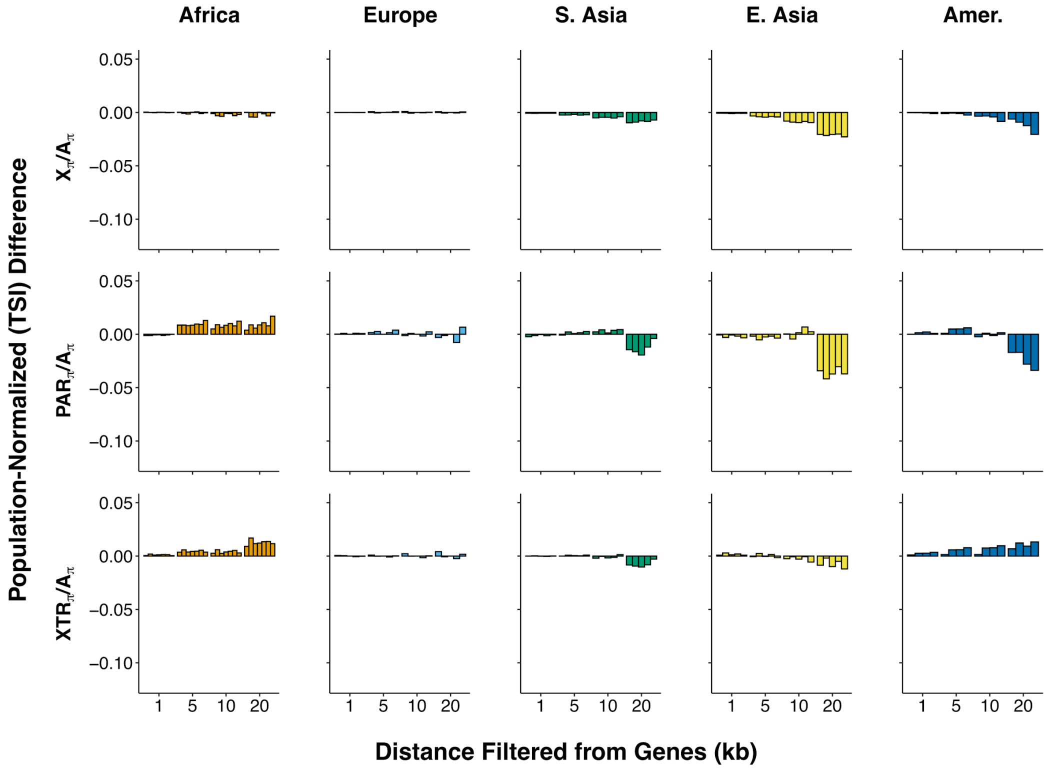
Africa
Europe
S. Asia
E. Asia
Amer.
Population-Normalized (TSI) Difference
$X_\pi/A_\pi$
$PAR_\pi/A_\pi$
$XTR_\pi/A_\pi$
0.05
0.00
−0.05
−0.10
1 5 10 20
Distance Filtered from Genes (kb)

b)

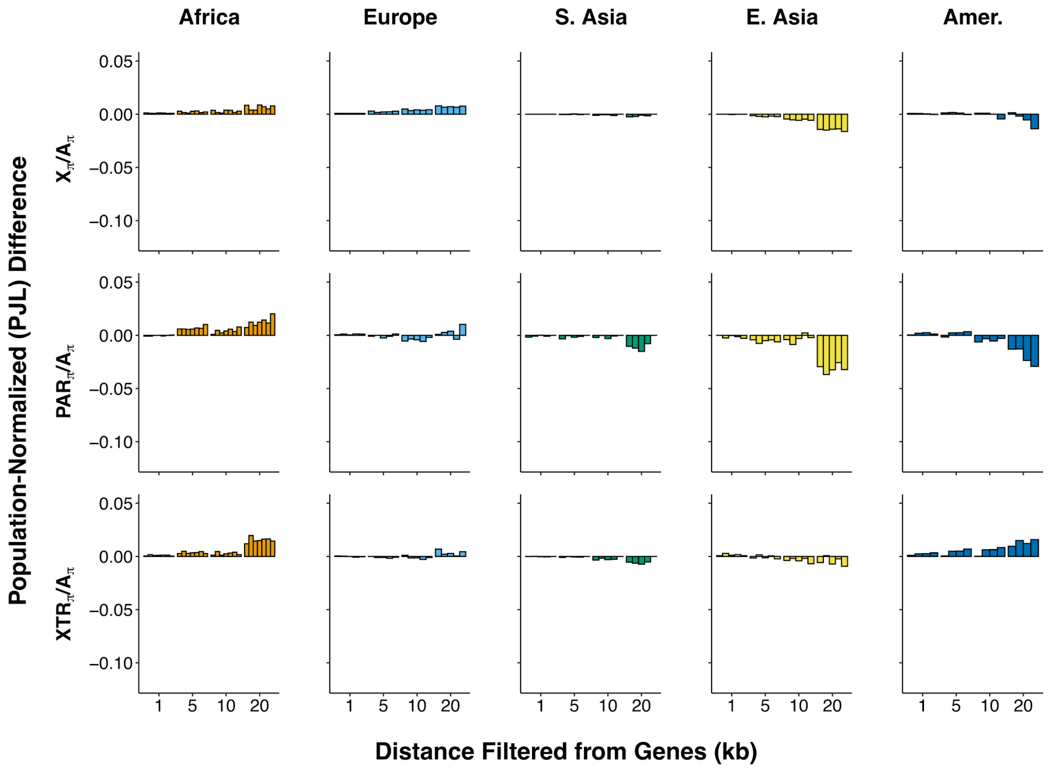
Africa
Europe
S. Asia
E. Asia
Amer.
Population-Normalized (PJL) Difference
$X_\pi/A_\pi$
$PAR_\pi/A_\pi$
$XTR_\pi/A_\pi$
0.05
0.00
−0.05
−0.10
1 5 10 20
Distance Filtered from Genes (kb)

c)

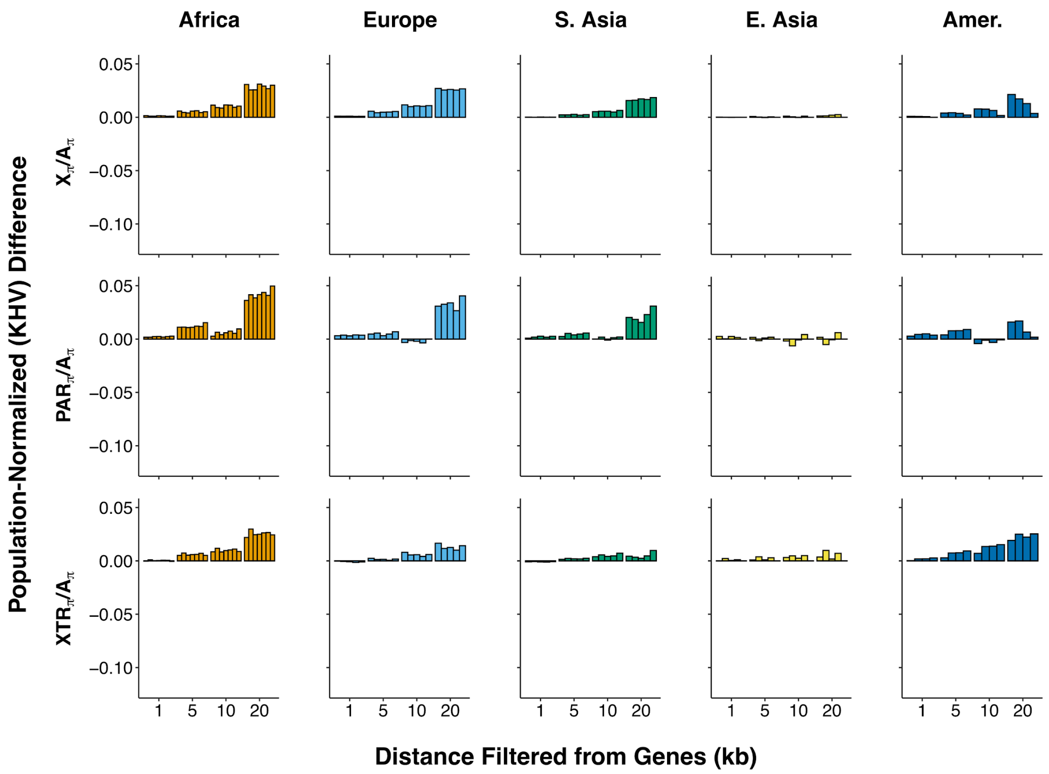


d)

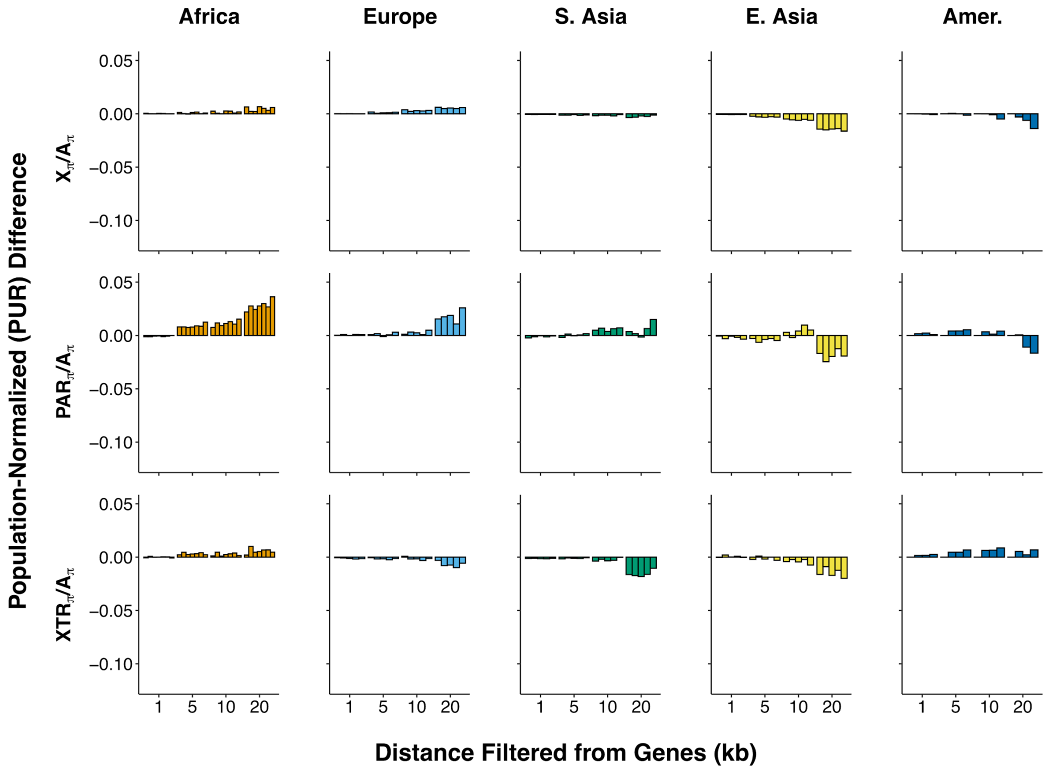


**Figure S3. Demography corrected ratios of X to autosomal diversity with increasing distance from genes across populations.** Diversity ratios between regions on the X chromosome—nonpseudoautosomal X (X), pseudoautosomal region 1 (PAR), and X-transposed region (XTR)—and autosomes for 25 1000 genomes populations. Values are reported as the difference between using a filter for only genes and a filter for 1kb, 5kb, 10kb, and 20kb from genes. These ratios are demography normalized by reporting each population relative to **A)** Toscani in Italia; **B)** Punjabi from Lahore, Pakistan; **C)** Kinh in Ho Chi Minh City, Vietnam; and **D)** Puerto Ricans from Puerto Rico. The order of populations is the same as reported in Figure 1B (less the corresponding population used for the correction).

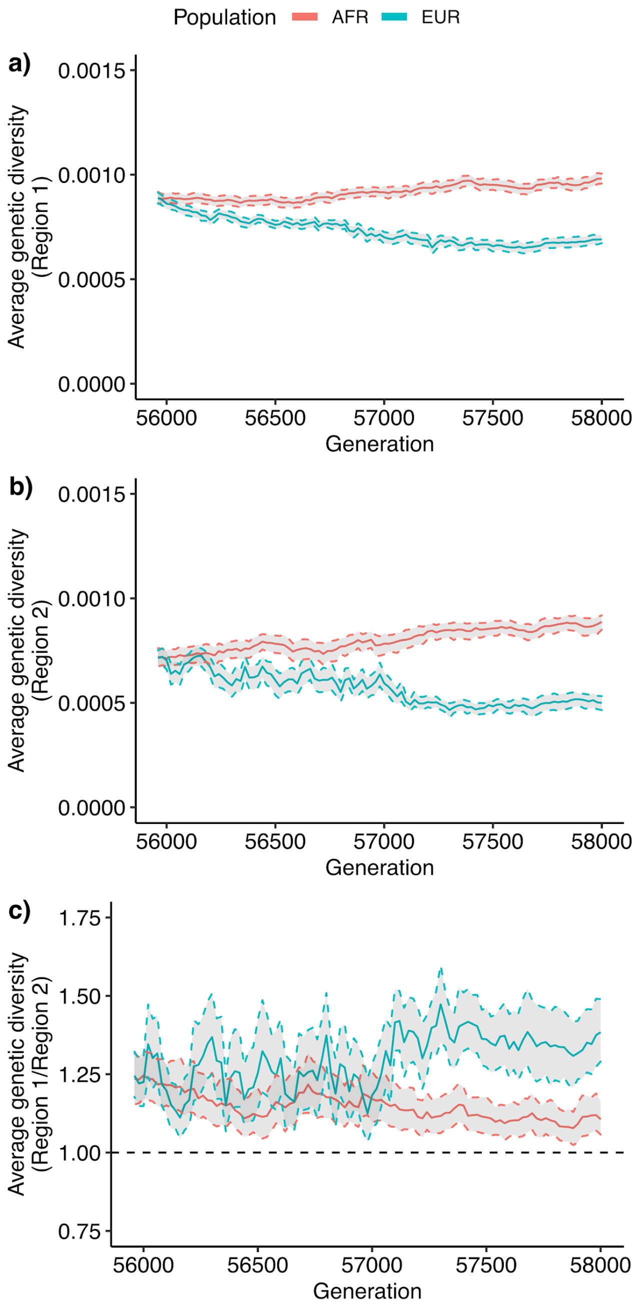


**Figure S4. Simulated genetic diversity in 2 regions with differing recombination rates under the Gravel et al. (2011) model of human evolution.** Diversity in two 50 kb regions with different recombination rates were tracked for 2000 generations following a European out-of-Africa bottleneck and expansion (83). Recombination rates were $2x10^{-7}$ in Region 1 and $1x10^{-8}$ in Region 2. Diversity, as calculated in Equation 1, is reported for Africans (red) and Europeans (blue) in **(A)** Region 1, **(B)** Region 2, and **(C)** after taking the Ratio of Region 1 to Region 2. Values are reported every 20 generations (from generation 55960 to 58000) by taking 250 samples of 100 individuals from each population and calculating the distribution of the diversity value in each region. The solid line is the mean and the dashed lines represent the 95% confidence intervals.

#### Supplementary Tables

**Table S1. 1000 Genomes populations used in analyses.** The number of male and female samples and population code used for each of the 26 1000 Genomes Project populations organized by superpopulation (African, Admixed American, East Asian, European, and South Asian)

| *Code* | *Description* | *Females* | *Males* | *Total* |
|---|---|---|---|---|
| **AFR** | **African** | **342** | **319** | **661** |
| ACB | African Caribbeans in Barbados | 49 | 47 | 96 |
| YRI | Yoruba in Ibadan, Nigeria | 56 | 52 | 108 |
| ASW | Americans of African Ancestry in SW USA | 35 | 26 | 61 |
| ESN | Esan in Nigeria | 46 | 53 | 99 |
| MSL | Mende in Sierra Leone | 43 | 42 | 85 |
| GWD | Gambian in Western Divisions in the Gambia | 58 | 55 | 113 |
| LWK | Luhya in Webuye, Kenya | 55 | 44 | 99 |
| **AMR** | **Admixed American** | **177** | **170** | **347** |
| MXL | Mexican Ancestry from Los Angeles USA | 32 | 32 | 64 |
| PUR | Puerto Ricans from Puerto Rico | 50 | 54 | 104 |
| CLM | Colombians from Medellin, Colombia | 51 | 43 | 94 |
| PEL | Peruvians from Lima, Peru | 44 | 41 | 85 |
| **EAS** | **East Asian** | **260** | **244** | **504** |
| CHB | Han Chinese in Beijing, China | 57 | 46 | 103 |
| JPT | Japanese in Tokyo, Japan | 48 | 56 | 104 |
| CHS | Southern Han Chinese | 53 | 52 | 105 |
| CDX | Chinese Dai in Xishuangbanna, China | 49 | 44 | 93 |
| KHV | Kinh in Ho Chi Minh City, Vietnam | 53 | 46 | 99 |
| **EUR** | **European** | **263** | **240** | **503** |
| CEU | Utah Residents (CEPH) with Northern and Western European Ancestry | 50 | 49 | 99 |
| TSI | Toscani in Italia | 54 | 53 | 107 |
| FIN | Finnish in Finland | 61 | 38 | 99 |
| GBR | British in England and Scotland | 45 | 46 | 91 |
| IBS | Iberian Population in Spain | 53 | 54 | 107 |
| **SAS** | **South Asian** | **229** | **260** | **489** |
| GIH | Gujarati Indian from Houston, Texas | 47 | 56 | 103 |
| PJL | Punjabi from Lahore, Pakistan | 48 | 48 | 96 |
| BEB | Bengali from Bangladesh | 44 | 42 | 86 |
| STU | Sri Lankan Tamil from the UK | 47 | 55 | 102 |
| ITU | Indian Telugu from the UK | 43 | 59 | 102 |

**Table S2. X chromosome diversity across populations.** Nucleotide diversity was calculated for each 1000 Genomes population and normalized for mutation rate using canFam3-hg19 divergence. P values are calculated using a permutation method with 10,000 replicates for the difference between a region (PAR1, XTR, or PAR2) and nonPAR. P-values here are not multiple-test corrected.

| Super Population | Population | Region | Diversity (pi) | P Value (vs nonPAR) |
|---|---|---|---|---|
| AFR | ACB | chr8 | 0.00361528 | ------ |
| AFR | ACB | nonPAR | 0.00251335 | ------ |
| AFR | ACB | PAR1 | 0.00441899 | < 0.0001 |
| AFR | ACB | PAR2 | 0.00377543 | 0.1233 |
| AFR | ACB | XTR | 0.00301991 | 0.0517 |
| AFR | ASW | chr8 | 0.00356146 | ------ |
| AFR | ASW | nonPAR | 0.00249919 | ------ |
| AFR | ASW | PAR1 | 0.00437048 | < 0.0001 |
| AFR | ASW | PAR2 | 0.00383831 | 0.0847 |
| AFR | ASW | XTR | 0.00295134 | 0.0582 |
| AFR | ESN | chr8 | 0.00357084 | ------ |
| AFR | ESN | nonPAR | 0.00247651 | ------ |
| AFR | ESN | PAR1 | 0.00442462 | < 0.0001 |
| AFR | ESN | PAR2 | 0.00379461 | 0.1068 |
| AFR | ESN | XTR | 0.00295609 | 0.0489 |
| AFR | GWD | chr8 | 0.00359202 | ------ |
| AFR | GWD | nonPAR | 0.0024744 | ------ |
| AFR | GWD | PAR1 | 0.00438228 | < 0.0001 |
| AFR | GWD | PAR2 | 0.00351761 | 0.1558 |
| AFR | GWD | XTR | 0.00299463 | 0.0378 |
| AFR | LWK | chr8 | 0.00360445 | ------ |
| AFR | LWK | nonPAR | 0.0025188 | ------ |
| AFR | LWK | PAR1 | 0.00433067 | < 0.0001 |
| AFR | LWK | PAR2 | 0.0038454 | 0.107 |
| AFR | LWK | XTR | 0.00303607 | 0.0143 |
| AFR | MSL | chr8 | 0.00362926 | ------ |
| AFR | MSL | nonPAR | 0.00253668 | ------ |
| AFR | MSL | PAR1 | 0.00446749 | < 0.0001 |
| AFR | MSL | PAR2 | 0.00378106 | 0.1241 |
| AFR | MSL | XTR | 0.0029912 | 0.0477 |
| AFR | YRI | chr8 | 0.00357637 | ------ |
| AFR | YRI | nonPAR | 0.00247075 | ------ |
| AFR | YRI | PAR1 | 0.00440654 | < 0.0001 |
| AFR | YRI | PAR2 | 0.0037438 | 0.1015 |
| AFR | YRI | XTR | 0.0029726 | 0.0408 |
| AMR | CLM | chr8 | 0.00281752 | ------ |
| AMR | CLM | nonPAR | 0.00174255 | ------ |
| AMR | CLM | PAR1 | 0.00348046 | < 0.0001 |
| AMR | CLM | PAR2 | 0.00294931 | 0.102 |

| | | | | |
|---|---|---|---|---|
| AMR | CLM | XTR | 0.00233712 | 0.0185 |
| AMR | MXL | chr8 | 0.00267508 | ------ |
| AMR | MXL | nonPAR | 0.00159974 | ------ |
| AMR | MXL | PAR1 | 0.00333288 | < 0.0001 |
| AMR | MXL | PAR2 | 0.00279026 | 0.0899 |
| AMR | MXL | XTR | 0.00226958 | 0.0137 |
| AMR | PEL | chr8 | 0.00248 | ------ |
| AMR | PEL | nonPAR | 0.00144861 | ------ |
| AMR | PEL | PAR1 | 0.00317044 | < 0.0001 |
| AMR | PEL | PAR2 | 0.00243977 | 0.1347 |
| AMR | PEL | XTR | 0.00194417 | 0.215 |
| AMR | PUR | chr8 | 0.00295736 | ------ |
| AMR | PUR | nonPAR | 0.00188871 | ------ |
| AMR | PUR | PAR1 | 0.00354182 | < 0.0001 |
| AMR | PUR | PAR2 | 0.00311649 | 0.0913 |
| AMR | PUR | XTR | 0.00237724 | 0.0292 |
| EAS | CDX | chr8 | 0.00251227 | ------ |
| EAS | CDX | nonPAR | 0.00140303 | ------ |
| EAS | CDX | PAR1 | 0.00327737 | < 0.0001 |
| EAS | CDX | PAR2 | 0.00244569 | 0.1338 |
| EAS | CDX | XTR | 0.0019594 | 0.0051 |
| EAS | CHB | chr8 | 0.00250122 | ------ |
| EAS | CHB | nonPAR | 0.00136881 | ------ |
| EAS | CHB | PAR1 | 0.00327971 | < 0.0001 |
| EAS | CHB | PAR2 | 0.00224832 | 0.1861 |
| EAS | CHB | XTR | 0.00195095 | 0.0033 |
| EAS | CHS | chr8 | 0.00249478 | ------ |
| EAS | CHS | nonPAR | 0.00137838 | ------ |
| EAS | CHS | PAR1 | 0.0032867 | < 0.0001 |
| EAS | CHS | PAR2 | 0.00237776 | 0.1464 |
| EAS | CHS | XTR | 0.00201036 | 0.0067 |
| EAS | JPT | chr8 | 0.00248974 | ------ |
| EAS | JPT | nonPAR | 0.00132271 | ------ |
| EAS | JPT | PAR1 | 0.00319639 | < 0.0001 |
| EAS | JPT | PAR2 | 0.00220501 | 0.2037 |
| EAS | JPT | XTR | 0.00204791 | 0.0011 |
| EAS | KHV | chr8 | 0.00251551 | ------ |
| EAS | KHV | nonPAR | 0.00140514 | ------ |
| EAS | KHV | PAR1 | 0.00335075 | < 0.0001 |
| EAS | KHV | PAR2 | 0.00241611 | 0.1336 |
| EAS | KHV | XTR | 0.00206352 | 0.0045 |
| EUR | CEU | chr8 | 0.0026069 | ------ |
| EUR | CEU | nonPAR | 0.0015599 | ------ |
| EUR | CEU | PAR1 | 0.00323275 | < 0.0001 |
| EUR | CEU | PAR2 | 0.00300167 | 0.0544 |
| EUR | CEU | XTR | 0.00214467 | 0.0317 |

| | | | | |
|---|---|---|---|---|
| EUR | FIN | chr8 | 0.00258571 | ------ |
| EUR | FIN | nonPAR | 0.00153039 | ------ |
| EUR | FIN | PAR1 | 0.00324371 | < 0.0001 |
| EUR | FIN | PAR2 | 0.00298957 | 0.058 |
| EUR | FIN | XTR | 0.00213324 | 0.0133 |
| EUR | GBR | chr8 | 0.00259937 | ------ |
| EUR | GBR | nonPAR | 0.00154916 | ------ |
| EUR | GBR | PAR1 | 0.0032688 | < 0.0001 |
| EUR | GBR | PAR2 | 0.00284047 | 0.0739 |
| EUR | GBR | XTR | 0.00215056 | 0.0169 |
| EUR | IBS | chr8 | 0.00263807 | ------ |
| EUR | IBS | nonPAR | 0.00157322 | ------ |
| EUR | IBS | PAR1 | 0.00328161 | < 0.0001 |
| EUR | IBS | PAR2 | 0.00283789 | 0.0849 |
| EUR | IBS | XTR | 0.00215956 | 0.0162 |
| EUR | TSI | chr8 | 0.00263909 | ------ |
| EUR | TSI | nonPAR | 0.00157384 | ------ |
| EUR | TSI | PAR1 | 0.00329901 | < 0.0001 |
| EUR | TSI | PAR2 | 0.00287421 | 0.0709 |
| EUR | TSI | XTR | 0.00221076 | 0.0133 |
| SAS | BEB | chr8 | 0.00274609 | ------ |
| SAS | BEB | nonPAR | 0.00161206 | ------ |
| SAS | BEB | PAR1 | 0.00348661 | < 0.0001 |
| SAS | BEB | PAR2 | 0.00311132 | 0.0641 |
| SAS | BEB | XTR | 0.0023667 | 0.0001 |
| SAS | GIH | chr8 | 0.00272361 | ------ |
| SAS | GIH | nonPAR | 0.00161177 | ------ |
| SAS | GIH | PAR1 | 0.00345347 | < 0.0001 |
| SAS | GIH | PAR2 | 0.00312773 | 0.0566 |
| SAS | GIH | XTR | 0.00234702 | 0.0022 |
| SAS | ITU | chr8 | 0.00274591 | ------ |
| SAS | ITU | nonPAR | 0.0016117 | ------ |
| SAS | ITU | PAR1 | 0.00349882 | < 0.0001 |
| SAS | ITU | PAR2 | 0.00318599 | 0.0489 |
| SAS | ITU | XTR | 0.00235697 | 0.0011 |
| SAS | PJL | chr8 | 0.00275393 | ------ |
| SAS | PJL | nonPAR | 0.00163613 | ------ |
| SAS | PJL | PAR1 | 0.0035451 | < 0.0001 |
| SAS | PJL | PAR2 | 0.00320998 | 0.0475 |
| SAS | PJL | XTR | 0.00232322 | 0.002 |
| SAS | STU | chr8 | 0.00275942 | ------ |
| SAS | STU | nonPAR | 0.00161187 | ------ |
| SAS | STU | PAR1 | 0.00352468 | < 0.0001 |
| SAS | STU | PAR2 | 0.00316304 | 0.0514 |
| SAS | STU | XTR | 0.00238356 | 0.0002 |

**Table S3. Filters with increasing distance from genes.** The amount of data remaining for each filter was calculated with increasing distance from genes (0kb, 1kb, 5kb. 10kb, 20kb, 50kb, and 100kb). Callable sites, variants, uncorrected diversity measures, and diversity corrected to canFam3 divergence are reported for each X chromosome region (PAR1, chrX, XTR, PAR2), the Y chromosome, and chromosome 8.

| | | Distance from Genes | | | | | | |
|---|---|---|---|---|---|---|---|---|
| | | **0 kb** | **1 kb** | **5 kb** | **10 kb** | **20 kb** | **50 kb** | **100 kb** |
| **PAR1** | **Callable Sites** | 439238 | 429019 | 395965 | 360584 | 286414 | 144245 | 24357 |
| | **Variant Sites** | 17742 | 17345 | 16076 | 14641 | 11602 | 5937 | 909 |
| | **Uncorrected Diversity** | 0.001885789 | 0.001889572 | 0.001911389 | 0.001900647 | 0.001925837 | 0.00202637 | 0.00165317 |
| | **Corrected Diversity (canFam3)** | 0.004376132 | 0.004390036 | 0.004432485 | 0.004406543 | 0.004462782 | 0.004727945 | 0.00359214 |
| **nonPAR** | **Callable Sites** | 61250366 | 59859475 | 55119179 | 50282402 | 42549990 | 28434242 | 15743791 |
| | **Variant Sites** | 1386828 | 1357783 | 1259724 | 1157551 | 991430 | 676655 | 384225 |
| | **Uncorrected Diversity** | 0.00070234 | 0.000705384 | 0.000714252 | 0.000722887 | 0.000736167 | 0.000753665 | 0.00077324 |
| | **Corrected Diversity (canFam3)** | 0.002404383 | 0.002415976 | 0.00244256 | 0.00247075 | 0.002508758 | 0.002554306 | 0.00258458 |
| **XTR** | **Callable Sites** | 1242053 | 1236090 | 1214578 | 1193863 | 1145085 | 992831 | 836805 |
| | **Variant Sites** | 29504 | 29354 | 28873 | 28437 | 27303 | 23790 | 20238 |
| | **Uncorrected Diversity** | 0.00092845 | 0.000923426 | 0.000925328 | 0.000924888 | 0.000920826 | 0.000928171 | 0.00093537 |
| | **Corrected Diversity (canFam3)** | 0.002999665 | 0.002966577 | 0.002944005 | 0.002972596 | 0.002704142 | 0.002793073 | 0.00289196 |
| **PAR2** | **Callable Sites** | 98048 | 95064 | 83750 | 66867 | 39255 | 0 | 0 |
| | **Variant Sites** | 3107 | 3022 | 2729 | 2200 | 1338 | 0 | 0 |
| | **Uncorrected Diversity** | 0.000928307 | 0.000945902 | 0.000990365 | 0.000983168 | 0.001098441 | NA | NA |
| | **Corrected Diversity (canFam3)** | 0.0034585 | 0.003573373 | 0.003836274 | 0.003743802 | 0.004119878 | NA | NA |
| **chr8** | **Callable Sites** | 49138677 | 47619394 | 42581683 | 37813522 | 30464701 | 17204730 | 7561320 |
| | **Variant Sites** | 1551553 | 1504169 | 1348229 | 1199442 | 967930 | 546508 | 239408 |
| | **Uncorrected Diversity** | 0.001106697 | 0.0011098 | 0.001118365 | 0.001122094 | 0.001122959 | 0.001119227 | 0.0011081 |
| | **Corrected Diversity (canFam3)** | 0.003513669 | 0.003526258 | 0.003558524 | 0.003576369 | 0.003590825 | 0.003592611 | 0.00357632 |

**Table S4. Gene density across the X chromosome and chromosome 8.** Gene length and the number of genes are reported for each region across the X chromosome (PAR1, PAR2, chrX, XTR) and chromosome 8.

| Region | Start | Stop | Region Length | Total Length of Genes | Fraction of Region that is Coding |
|---|---|---|---|---|---|
| **chr8** | 1 | 146364022 | 146364022 | 79534155 | 0.54339963 |
| **nonPAR**[1] | 2699520 | 88193855 | 147231524 | 58653704 | 0.39837735 |
| | 93193855 | 154931044 | - | - | - |
| **PAR1**[2] | 60001 | 2699520 | 2639519 | 985328 | 0.37329832 |
| **PAR2**[2] | 154931044 | 155260560 | 329516 | 174171 | 0.52856614 |
| **XTR**[3] | 88193855 | 93193855 | 5000000 | 934266 | 0.1868532 |

1. nonPAR is calculated as the remaining regions after removing PAR1, PAR2 and XTR. Thus it is split up into two non-contiguous regions.
2. PAR1 and PAR2 coordinates come from the hg19 region definitions.
3. XTR is defined between 88 and 93 Mb (35). It consists of two homologous blocks within this region and between the Y chromosome. We use these coordinates to be as conservative as possible.
